# Physiological robustness of MScanFit to simulated motor unit loss and remodelling

**DOI:** 10.64898/2026.08.28.747933

**Authors:** Dawood Ahmed, Mohammed Almokdad, Kelvin E. Jones

**Affiliations:** Department of Physiology, University of Alberta, Edmonton, AB, Canada; Neuroscience and Mental Health Institute, University of Alberta, Edmonton, AB, Canada; Faculty of Kinesiology, Sport, & Recreation, University of Alberta, Edmonton, AB, Canada

## Abstract

**Introduction:** MScanFit estimates motor unit number and size from compound muscle action potential (CMAP) scans, but its accuracy may depend on the physiological processes that shape motor unit loss and remodelling. We reproduced the general dynamic remodelling framework used in the original MScanFit simulation and expanded it to test MScanFit robustness across physiologically heterogeneous conditions.

**Methods:** Generative computational models simulated CMAP scans from motor unit pools undergoing progressive loss from 160 to 5 surviving motor units. Twelve remodelling conditions combined denervation pattern (random or selective), reinnervation method (random, distributive, or size-weighted), and neuromuscular resilience (20% or 60%); two additional conditions modelled denervation without collateral reinnervation. MScanFit estimates were compared with the known motor unit numbers and mean motor unit sizes used to generate each scan.

**Results:** MScanFit reproduced the principal behaviour reported in the original simulation and generally tracked progressive motor unit loss across heterogeneous remodelling conditions. However, motor unit number estimation (MUNE) error changed systematically with remodelling physiology. Selective denervation and distributive reinnervation increased error at several intermediate stages of motor unit loss, whereas 60% resilience produced greater error than 20% resilience at every stage after remodelling began. Motor unit size estimates also tracked the underlying increase in mean unit size, but their accuracy was more strongly affected by remodelling, particularly with 60% resilience and advanced motor unit loss.

**Conclusion:** MScanFit motor unit number estimates were broadly robust to substantial motor unit loss and neuromuscular remodelling but were not physiologically invariant. Denervation and collateral reinnervation systematically influenced the magnitude and direction of estimation error, indicating that variation in MScanFit performance can arise from differences in the underlying motor unit population. These findings support MScanFit as a robust measure of motor unit loss across heterogeneous neuromuscular phenotypes and show that some of its estimation variability has a physiological basis.

## 1. Introduction

A motor unit comprises an α-motor neuron and the skeletal muscle fibres it innervates. ^1,2^ Motor units differ substantially in their contractile and electrophysiological properties. Innervation ratio, the number of fibres innervated by an individual motor neuron, contributes to both motor unit force and the size of its surface-recorded potential. These differences are commonly represented by slow, fast fatigue - resistant, and fast fatigable motor unit classes as established in experimental preparations. ^1–5^ Larger, faster motor units generally have greater innervation ratios and produce larger surface-recorded potentials than smaller, slower units. Thus, the composition of a motor unit pool can influence the measurements used to estimate motor unit number.

Motor neuron degeneration changes both the number and organization of functioning motor units. In amyotrophic lateral sclerosis (ALS), progressive motor neuron loss denervates skeletal muscle fibres, but surviving motor axons may compensate and reinnervate via collateral sprouting, enlarging the innervation ratios of these surviving units. Denervation and reinnervation are not necessarily random. Experimental models show the preferential vulnerability of larger, fast motor units, whereas slower units are relatively resistant and may retain greater capacity for collateral sprouting. ^6–11^ Consequently, motor unit pools containing the same number of surviving units may differ substantially in their size distribution and in how denervated muscle fibres are redistributed among survivors. Such differences could influence electrophysiological estimates of motor unit numbers independently of the true number remaining.

Direct counting of functioning motor units is not possible in intact human muscle, so motor unit number estimation (MUNE) methods instead infer their number from electrophysiological measurements. ^12,13^ Several newer approaches use the compound muscle action potential (CMAP) scan, which records the progressive growth of the muscle response as stimulus intensity is adjusted monotonically in small increments. ^14–18^ MScanFit fits a model of motor unit recruitment to the CMAP scan and estimates both the number of motor units and their underlying size distribution. ^19–22^ Its accuracy therefore depends on how well the fitted model represents the motor unit population that generated the scan.

As part of the original MScanFit report, Bostock used simulated CMAP scans with known motor unit populations to demonstrate the performance of the fitting procedure under a simplified model of progressive motor unit loss and collateral reinnervation. ^19^ Generative models are particularly useful for algorithm testing because the number and sizes of the motor units producing each simulated CMAP scan are known, unlike in vivo. ^23–26^ However, performance under one set of remodelling assumptions does not establish that an algorithm will behave similarly when denervation and collateral reinnervation alter the motor unit population in different ways. The selective loss of larger units, alternative distributions of recovered innervation among surviving units, and differences in collateral reinnervation capacity could each change CMAP scan morphology and the resulting MScanFit estimate.

The present study reproduced the general dynamic remodelling framework used in Bostock’s original MScanFit simulation, retaining the same sequence of motor unit pool sizes (160, 80, 40, 20, 10, and 5) to facilitate benchmarking against published results.^19^ We then expanded the remodelling space by varying denervation pattern, collateral reinnervation method, and neuromuscular resilience, which controlled the extent of collateral reinnervation. The primary objective was to determine whether MScanFit (referring to MSF 2, the latest and default algorithm, unless otherwise specified) motor unit number estimates remained reasonably accurate across these heterogeneous remodelling conditions and stages of motor unit loss. We distinguished robustness from invariance by also examining whether the magnitude and direction of MUNE error changed systematically with the underlying remodelling physiology. Secondary analyses examined the accuracy of MScanFit estimates of mean motor unit size.

## 2. Methods

### 2.1 OVERVIEW

A generative computational model was used to evaluate MScanFit so that the motor unit number and SMUP amplitudes underlying each simulated CMAP scan would be known. The model operated at the level of the motor unit rather than individual muscle fibres. Each motor unit was characterized by its single motor unit potential (SMUP) amplitude, mean activation threshold, and relative threshold spread, which together determined its contribution to the simulated CMAP scan.

Model development proceeded in two stages. First, we implemented the general dynamic remodelling framework used in the original MScanFit (MSF1) simulation to establish a benchmark against published results.^19^ Selected parameters were then calibrated against experimental CMAP scans recorded from the abductor pollicis brevis (APB) muscle of healthy participants. The calibrated model was used for the principal simulation experiment.

Fourteen neuromuscular remodelling conditions were evaluated across six stages of motor unit loss (160, 80, 40, 20, 10, and 5 surviving units). Thirty stochastic trajectories, representing thirty simulated individuals, were generated for each condition, with remodelling retained between stages to produce path-dependent histories. A simulated CMAP scan was generated and analysed with MScanFit at each stage, producing 2,520 scans in total. MScanFit estimates were compared with the known motor unit number and SMUP amplitudes used to generate each scan.

### 2.2 INITIAL MOTOR UNIT POOLS

Each simulated motor unit pool initially contained 160 motor units, representing the upper range of published MScanFit estimates for the abductor pollicis brevis (APB) muscle. ^22^ The APB was selected because it is commonly used for CMAP scan MUNE and can be activated reliably by median nerve stimulation.

Each motor unit *i* was defined by three properties: SMUP amplitude (*µ*_*i*_ ), mean activation threshold (*τ*_*i*_ ), and relative spread (*ρ*).^15,19^ SMUP amplitude represented a unit’s contribution to the surface-recorded electromyographic response to nerve stimulation. Individual muscle fibres, motor unit potential waveforms, and spatial relationships within the muscle were not modelled explicitly. SMUP amplitudes were sampled independently from a shifted exponential distribution,

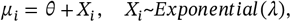

where *θ* defines the minimum SMUP amplitude and *λ* determines the distribution above this minimum. This distribution produced many small units and progressively fewer large units (Figure 1), consistent with the positively skewed distribution of experimental surface-recorded SMUP amplitudes for the APB muscle. ^27,28^

**Figure 1.**
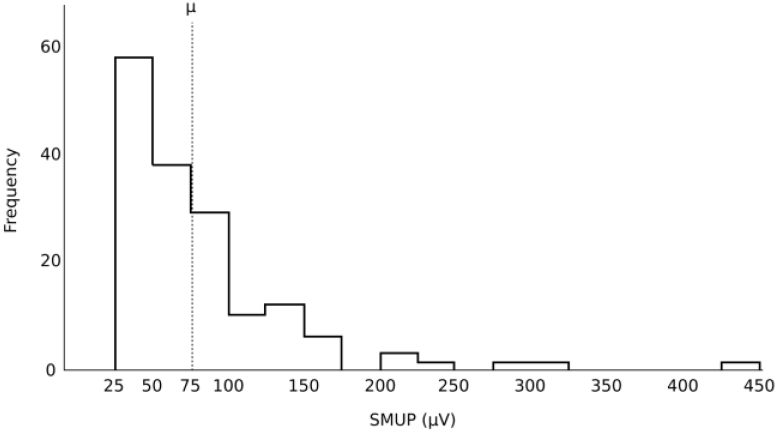
Distribution of SMUP amplitudes in an initial simulated motor unit pool. Representative frequency distribution of SMUP amplitudes in an initial abductor pollicis brevis (APB) motor unit pool (N = 160). SMUP amplitudes were drawn from a shifted exponential distribution, producing many small units and progressively fewer large units. No SMUP amplitudes were below the imposed 25 *µ*V minimum. A dotted line indicates the mean (*µ*) SMUP amplitude of the population.

Mean activation thresholds were sampled independently of SMUP amplitude from a Gaussian distribution,

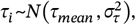

reflecting the approximately random order of motor axon activation during percutaneous electrical nerve stimulation. Unlike voluntary motor unit recruitment, activation during external stimulation depends on both axonal excitability and the position of individual axons within the applied electrical field. ^25^

Activation of each motor axon was probabilistic rather than occurring at a fixed stimulus intensity. Threshold variability was specified by a relative threshold spread of *ρ* = 1.65%, consistent with previous human axonal excitability measurements.^29^ The resulting activation probability was used to generate CMAP scans as described in Section 2.3.

Thirty independent stochastic realizations of the initial 160 - unit pool were generated for each simulation set. These differed in their sampled SMUP amplitudes and activation thresholds. Distribution parameters used in the principal simulations were determined during model calibration and are reported in Section 2.4.

### 2.3 SIMULATING CMAP SCANS

Once a motor unit pool was generated, a corresponding CMAP scan was simulated by calculating the probabilistic activation of each unit across increasing stimulus intensities. Each scan contained 500 stimulus intensities geometrically spaced from 0.5 mA below the lowest mean activation threshold in the pool to 0.5 mA above the highest mean activation threshold. ^19,30^ Twenty constant stimuli were added at each boundary to provide the pre- and post-scan regions used during MScanFit analysis.

For a motor unit *n*, its activation threshold varied around its mean threshold *τ*_*n*_ . The standard deviation of this threshold was derived from the relative threshold spread *ρ*:

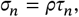

where *ρ* = 0.0165 . At stimulus intensity *x*, the probability that unit *n* was activated was given by the cumulative distribution function of a Gaussian distribution (*CDF*_*G*_),

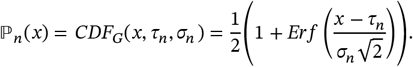

Activation was determined stochastically at each stimulus intensity. For each motor unit and stimulus intensity, *U*_*n,x*_ was sampled independently from a uniform distribution, *U*(0,1), and the unit was considered active when:

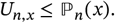

Because motor units were represented by SMUP amplitude rather than an explicitly modelled waveform, activation contributed the full SMUP amplitude, *µ*_*n*_, to the simulated response. The noise-free CMAP amplitude at stimulus intensity x was therefore:

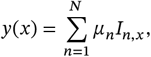

where *I*_*n,x*_ = 1 when motor unit *n* was activated and *I*_*n,x*_ = 0 otherwise. Independent additive recording noise was then applied to each simulated CMAP response, including the 20 pre-scan and post-scan points :

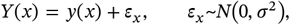

where *σ* = 10 *μ*V. Thus, repeated scans generated from the same underlying motor unit pool could differ because both motor unit activation and recording noise were stochastic, whereas motor unit number, SMUP amplitudes, and mean activation thresholds remained fixed.

### 2.4 CMAP SCAN CALIBRATION

The generative model was first implemented using parameters adopted from the original MScanFit simulation. ^19^ Initial pools contained 160 motor units, with SMUP amplitudes sampled from a shifted exponential distribution with a minimum of 25 *μ*V and a mean of 62.5 *μ*V. Mean activation thresholds were sampled independently from a Gaussian distribution with a mean of 26.5 mA and standard deviation of 2.0 mA. These parameters were used to establish that our independent implementation generated CMAP scans and MScanFit estimates comparable to those reported by Bostock before calibration for the principal simulations.

Selected model parameters were then calibrated against experimental APB CMAP scans from healthy participants. These recordings were obtained from a previously published multicentre study of MScanFit repeatability for which the acquisition procedures have been described in detail. ^22^ The present analysis included all 38 APB CMAP scans available from sites covered by an existing secondary analysis agreement; no additional selection or quality-control criteria were applied.

Calibration targeted five characteristics of experimental CMAP scans: the maximal CMAP amplitude; the stimulus intensities corresponding to 5%, 50%, and 95% of maximal CMAP amplitude (S5, S50, and S95); and the slope of the stimulus–response relationship at S50. Three model parameters were optimized: the mean SMUP amplitude, mean activation threshold, and the standard deviation of the activation-threshold distribution. The minimum SMUP amplitude and relative threshold spread remained fixed at 25 *μ*V and 1.65%, respectively.

Parameter values were estimated using Gaussian process - based Bayesian optimization. ^31^ For each candidate parameter set, 30 independent stochastic realizations were generated, and the five calibration features were calculated for each simulated CMAP scan. The mean of each feature across the 30 simulations was compared with the corresponding mean from the experimental scans. The objective function was the sum of squared relative differences between simulated and experimental feature means:

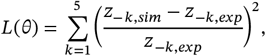

where *θ* denotes the candidate parameter set and *z*_−*k,sim*_ and *z*_−*k,exp*_ are the simulated and experimental means for calibration feature *k*. Using 30 realizations for each candidate parameter set reduced the influence of random variation on the optimization. The parameter set minimizing the objective function *L*(*θ*) was retained for the principal simulations (Figure 2).

**Figure 2.**
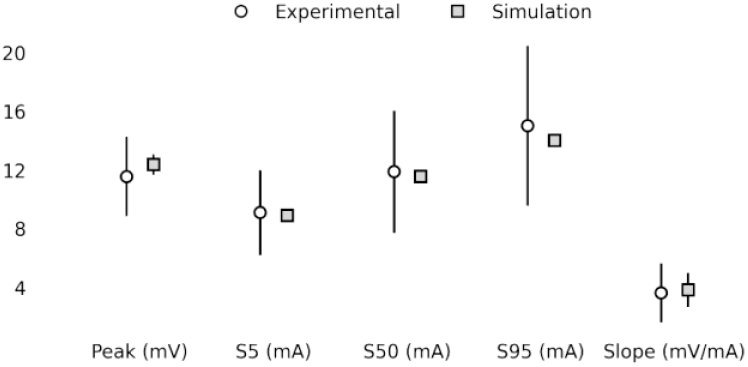
Calibration of simulated CMAP scans to experimental APB recordings. Comparison of peak CMAP amplitude, S5, S50, S95, and midpoint slope between experimental (n = 38) and simulated (n = 30) CMAP scans of the abductor pollicis brevis (APB) muscle. Symbols indicate means and vertical bars indicate standard deviation.

Calibration produced a mean SMUP amplitude of 75.0 *μ*V and an activation threshold distribution with a mean of 11.9 mA and standard deviation of 1.5 2 mA. These values were held constant as the initial distribution parameters for all subsequent neuromuscular remodelling simulations.

### 2.5 NEUROMUSCULAR REMODEL L ING

Neuromuscular remodelling was simulated by progressively reducing each motor unit pool from 160 to 80, 40, 20, 10, and 5 surviving units. At each transition, half of the motor units present at the preceding stage were removed according to one of two denervation rules. Under **random denervation**, units were selected for removal at random, as in the original MScanFit simulation. ^19^ Under **selective denervation**, units with the largest SMUP amplitudes were removed deterministically . Because SMUP amplitudes could be increased by collateral reinnervation and these changes were retained between stages, selective denervation depended on the accumulated remodelling history of each trajectory.

Collateral reinnervation was represented at the level of SMUP amplitude because individual muscle fibres were not modelled explicitly. Let *D* denote the set of motor units removed during a denervation step and *µ*_*j*_ the SMUP amplitude of denervated motor unit *j*. The total SMUP amplitude associated with the denervated units was:

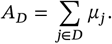

Neuromuscular resilience, *r*, was the model parameter controlling the extent of collateral reinnervation. For the principal remodelling conditions, *r* = 0.20 or 0.60; two additional boundary conditions used *r* = 0, representing denervation without collateral reinnervation. The implementation of resilience differed according to the reinnervation rule. Under **random reinnervation**, adapted from the original model, ^19^ *r* specified the proportion of denervated motor units selected for reinnervation. The entire SMUP amplitude of each selected unit was transferred to a randomly selected surviving motor unit, with each surviving motor unit receiving the amplitude of no more than one denervated motor unit during a remodelling step.

Under **distributive reinnervation**, the total recovered amplitude, *A*_*R*_ = *rA*_*D*_, was divided equally among all surviving motor units in set *S*:

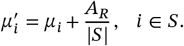

Under **size-weighted reinnervation**, recovered amplitude was distributed among surviving units in proportion to the square of their current SMUP amplitudes:

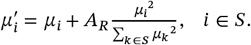

This method was motivated by the size-dependent collateral reinnervation represented in the dynamic muscle model of Sleutjes,^26^ such that larger surviving motor units received a disproportionately greater share of the recovered amplitude.

For a given set of denervated units and resilience level, distributive and size-weighted reinnervation preserved the same total recovered SMUP amplitude and differed only in how that amplitude was distributed among surviving units. Under random reinnervation, the recovered amplitude depended on which denervated motor units were selected for complete amplitude transfer. Reinnervation increased the SMUP amplitudes of surviving units but did not alter motor unit number. Updated SMUP amplitudes were retained at subsequent stages, producing cumulative, path - dependent remodelling within each trajectory.

The principal experimental design comprised two denervation rules, three reinnervation rules, and two resilience levels (20% and 60%), forming a balanced 2 × 3 × 2 factorial design with 12 remodelling conditions. Two additional no-reinnervation conditions combined random or selective denervation with *r* = 0; because no amplitude was recovered, the reinnervation rule was undefined. In total, 14 simulation conditions were evaluated across six motor unit stages and 30 stochastic trajectories per condition (Table 1 ).

**Table 1.** Overview of 14 neuromuscular remodelling experimental conditions.

| Condition | Denervation | Reinnervation | Resilience | Code |
|---|---|---|---|---|
| 1 | random | random | 20% | RR20 |
| 2 | random | random | 60% | RR60 |
| 3 | random | distributive | 20% | RD20 |
| 4 | random | distributive | 60% | RD60 |
| 5 | random | size-weighted | 20% | RS20 |
| 6 | random | size-weighted | 60% | RS60 |
| 7 | random | none | - | RNone |
| 8 | selective | random | 20% | SR20 |
| 9 | selective | random | 60% | SR60 |
| 10 | selective | distributive | 20% | SD20 |
| 11 | selective | distributive | 60% | SD60 |
| 12 | selective | size-weighted | 20% | SS20 |
| 13 | selective | size-weighted | 60% | SS60 |
| 14 | selective | none | - | SNone |

### 2.6 MScanFit ANALYSIS

Each simulated CMAP scan was analysed using MScanFit within the QTracW software suite (© University College London, UK; distributed by Digitimer Ltd, Welwyn Garden City, UK). The default MScanFit fitting procedure (MSF2) was used with default settings except that the pre-scan and post-scan regions were fixed to the first and last 20 stimulus points, respectively, corresponding to the constant stimuli incorporated during CMAP scan generation. The minimum motor unit amplitude was set to 25 *μ*V, matching the lower bound of the SMUP amplitude distribution used to generate the simulated motor unit pools. For each scan, MScanFit motor unit number estimate (MUNE) and mean single-unit estimate (MSUE) were retained for comparison with the known motor unit number and mean SMUP amplitude of the simulated pool . For the benchmark comparison (Section 3), RR60 scans were additionally fitted with MSF1 to match Bostock’s original procedure.

### 2.7 OUTCOME MEASURES

The primary outcome was the accuracy of the MScanFit motor unit number estimate (MUNE) relative to the known number of motor units in each simulated pool. Signed percentage error quantified the direction and magnitude of estimation bias:

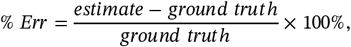

where negative values indicate underestimation and positive values indicate overestimation. Absolute percentage error quantified estimation accuracy irrespective of direction:

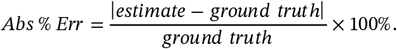

MUNE absolute percentage error was the primary outcome used for comparisons among remodelling conditions. The secondary outcome was the accuracy of motor unit size estimates. The target mean SMUP amplitude was calculated directly from the simulated motor unit population and compared with the mean single unit estimate (MSUE) generated by MScanFit. Signed error (Err; *μ*V) quantified the direction of estimation bias, whereas absolute error (Abs Err; *μ*V) was used as the primary measure of MSUE accuracy.

### 2.8 DATA PROCESSING AND ANALYSIS

The statistical analyses of neuromuscular remodelling focused on the 12 conditions forming a balanced 2 × 3 × 2 factorial design: denervation mechanism (random, selective), reinnervation mechanism (random, distributive, size-weighted), and resilience (20%, 60%). Each condition comprised 30 independent trajectories followed across six stages of motor unit loss (160, 80, 40, 20, 10, and 5 motor units). The two conditions without collateral reinnervation (*RNone* and *SNone*) were analysed separately.

An estimation-statistics framework was used to emphasize effect magnitude and precision. 32,33 At each motor unit stage, marginal mean differences were estimated for selective versus random denervation, distributive versus random reinnervation, size-weighted versus random reinnervation, and 60% versus 20% resilience, averaging across the balanced levels of the other factors. Bias-corrected and accelerated (BCa) 95% confidence intervals were obtained from 9,999 bootstrap resamples, using trajectory as the resampling unit and resampling independently within simulation condition. The 160-motor unit stage, before remodelling occurred, was retained as an internal control.

This stage-specific analysis was applied separately to MUNE absolute percentage error and MSUE absolute error. The no - reinnervation conditions were compared by estimating the stage-specific mean difference between *SNone* and *RNone*. Comparisons with the original MScanFit simulation reported by Bostock were descriptive because the underlying simulation data were unavailable. 19 Analyses were performed in Python using Pandas, NumPy, and SciPy, with figures generated using Matplotlib. The complete simulated dataset and MScanFit outputs used for these analyses are publicly available on Figshare (doi:10.6084/m9.figshare.33190827).

## 3. Results

Results are presented in five parts. First, the *RR60* condition was benchmarked against Bostock’s original MScanFit (MSF1) simulation. Second, we confirmed that the expanded remodelling rules generated heterogeneous motor unit populations and CMAP trajectories. Third, we evaluated MScanFit motor unit number accuracy across conditions and stages of motor unit loss. remodelling Fourth, we examined the accuracy of MScanFit estimates of mean motor unit size. Finally, we examined the accuracy of MScanFit when collateral reinnervation was absent. MScanFit tracked progressive motor unit loss across all remodelling conditions, but its accuracy depended on the underlying physiology . Size estimates followed the rise in mean unit size but degraded more sharply with remodelling .

### BENCHMARK COMPARISON WITH ORIGINAL MScanFit

#### SIMULATION

The *RR60* condition was designed to approximate the simulation used in Bostock’s original evaluation of MSF1,^19^ using comparable random motor unit loss and collateral reinnervation, but with 30 rather than 10 simulated trajectories. Representative CMAP scans from two *RR60* trajectories illustrate the progressive changes in scan morphology produced by motor unit loss and remodelling (Figure 3). Table 2 summarizes MS F1 performance across the 180 resulting *RR60* scans .

**Table 2.** Results of fitting 180 simulated CMAP scans from the RR60 condition with MScanFit (MSF1).

| True Motor Units (n) | 5 (30) | 10 (30) | 20 (30) | 40 (30) | 80 (30) | 160 (30) | Combined (180) |
|---|---|---|---|---|---|---|---|
| MUNE | 5.2 ± 0.4 | 10.9 ± 1.1 | 22.8 ± 3.8 | 41.9 ± 4.1 | 83.4 ± 8.2 | 148.4 ± 14.7 | — |
| MUNE % Err | 4.7 ± 8.6 | 9.0 ± 10.9 | 14.0 ± 18.9 | 4.7 ± 10.2 | 4.2 ± 10.2 | −7.2 ± 9.2 | 4.9 ± 13.3 |
| MUNE Abs % Err | 4.7 ± 8.6 | 11.0 ± 8.8 | 17.0 ± 16.1 | 8.8 ± 6.8 | 8.2 ± 7.3 | 8.9 ± 7.5 | 9.8 ± 10.3 |
| MSUE (µV) | 738.5 ± 123.0 | 454.4 ± 68.1 | 278.4 ± 48.9 | 186.4 ± 20.6 | 117.7 ± 14.6 | 83.1 ± 11.4 | — |
| MSUE Abs Err (µV) | 31.0 ± 54.2 | 46.6 ± 32.4 | 41.5 ± 33.1 | 15.9 ± 11.5 | 9.4 ± 7.7 | 8.1 ± 8.1 | 25.4 ± 33.1 |

*Values indicate mean ± SD. MUNE: motor unit number estimate; Abs: absolute value; MSUE: mean single-unit estimate.*

**Figure 3.**
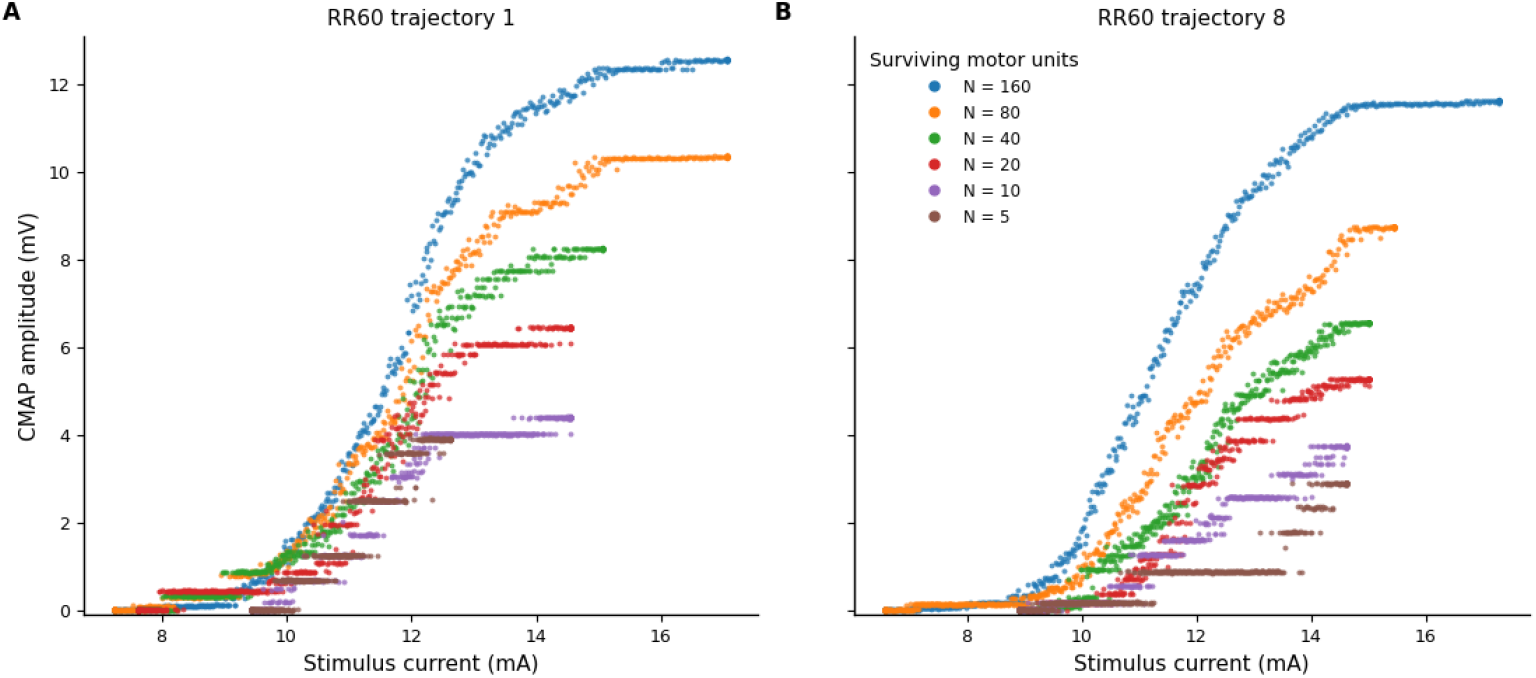
Representative simulated CMAP scans from two RR60 trajectories across progressive motor unit loss. CMAP scans are shown for two of the 30 simulated trajectories generated under the RR60 remodelling condition. Within each panel, scans from six stages of degeneration are superimposed (*N* = 160, 80, 40, 20, 10, and 5 surviving motor units), with CMAP amplitude plotted against stimulus current. Each point represents the simulated response to a single stimulus in the scan sequence. As motor unit loss progresses, the maximum CMAP decreases and the scan morphology becomes increasingly discontinuous, with larger step-like increments becoming especially apparent at *N* = 20 and below. These steps reflect the recruitment of progressively fewer, relatively larger motor units. The two trajectories also illustrate that, even within the same remodelling condition, the pattern and magnitude of CMAP decline are not identical. For example, from *N* = 160 to *N* = 80, maximum CMAP decreased by 2.23 mV in trajectory 1 (12.58 to 10.36 mV) and by 2.89 mV in trajectory 8 (11.63 to 8.75 mV). Together, these examples illustrate both the expected effects of degeneration on CMAP scans and the trajectory-to-trajectory variability present in the simulations.

MSF1 estimates generally tracked the true motor unit number in both the RR60 and Bostock simulations across the full range of motor unit pool sizes (Figure 4A). Absolute percentage errors were larger for RR60 at 5 –20 motor units, most notably at 10 and 20 unit s, but were comparable or smaller at 40 –160 units (Figure 4C). Percentage error is amplified at small integer pool sizes, where an error of one unit corresponds to 20% at 5 units and 10% at 10 units. Overall MUNE absolute percentage error was 9.8 ± 10.3% for RR60 compared with 6.9 ± 7.8% reported by Bostock. Mean SMUP size increased more rapidly with motor unit loss in RR60 than in Bostock’s simulation (Figure 4B). MSF1 generally followed these differing target distributions, although MSUE absolute error was greater in RR60 at 5 20 motor units, peaking at 10 –20 units (Figure 4D).

**Figure 4.**
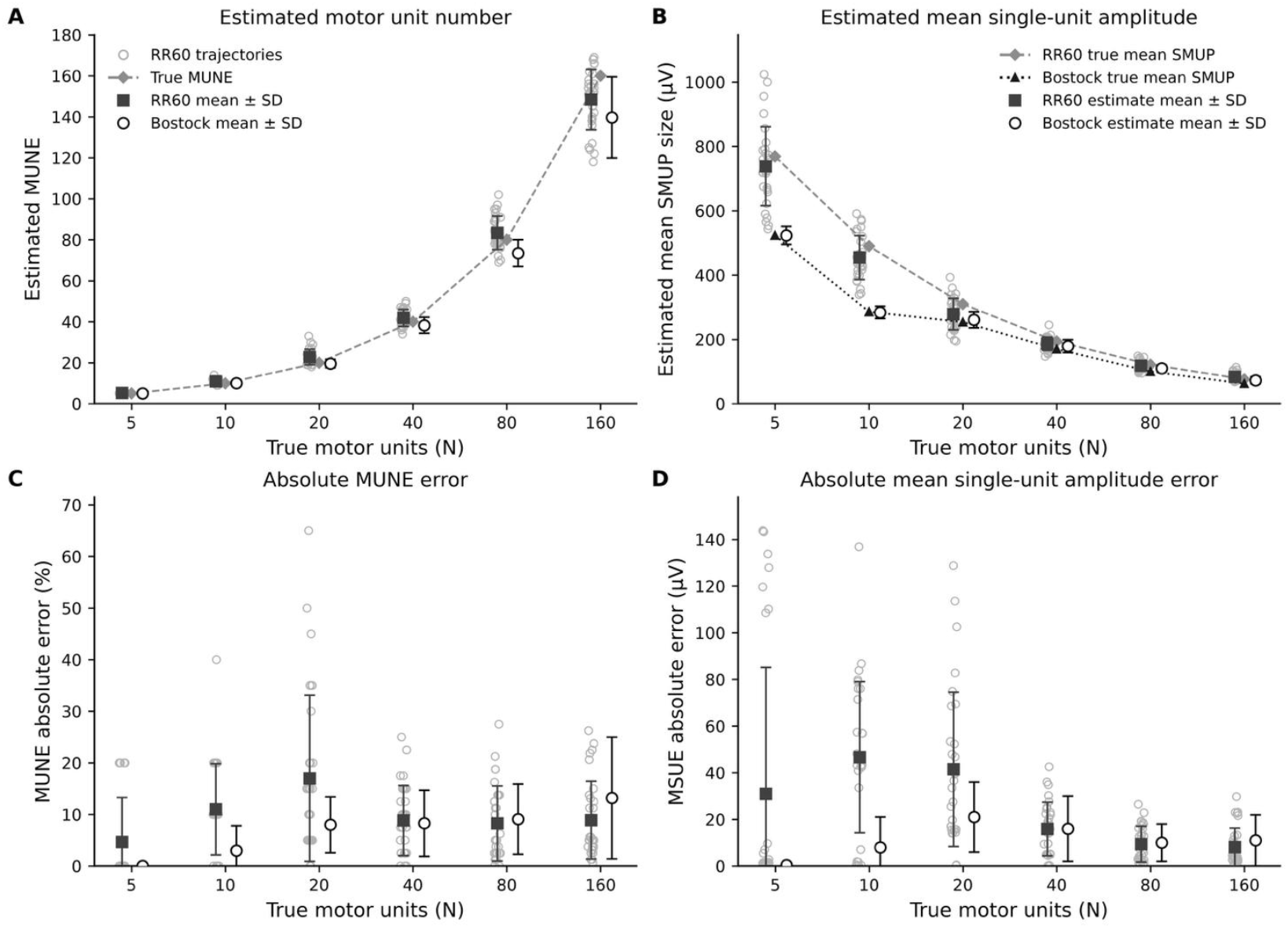
Benchmark comparison of MScanFit (MSF1) performanceon two generative models. (A) MUNE estimates and (B) estimates of mean single unit amplitude (MSUE) across true motor pool sizes. Individual RR60 results are shown for 30 simulated trajectories at each pool size, with mean ± SD. Bostock values are published results from 10 simulated scans per pool size. Reference lines in (A) indicate the true motor unit numbers. Reference lines in (B) show the target mean SMUP amplitudes for the RR60 and Bostock simulations. (C) MUNE absolute percentage error and (D) MSUE absolute error for the same simulations. RR60 and Bostock summary values were offset slightly within each motor unit category to prevent overlap; the horizontal offset is for visual clarity only and has no analytical meaning . No inferential statistical comparisons were performed.

### SIMULATED NEUROMUSCULAR REMODEL L ING PRODUCED HETEROGENEOUS MOTOR UNIT POPULATIONS

Before evaluating MScanFit performance, we examined whether the remodelling rules produced meaningfully different motor unit populations. At the initial stage (*N* = 160), all conditions were generated from the same underlying distributions of SMUP size and activation threshold and therefore began from comparable motor unit populations. As degeneration progressed, the simulations diverged in their whole-muscle behaviour. Maximum CMAP declined in all conditions, but the magnitude and trajectory of decline differed according to denervation, reinnervation, and resilience (Figure 5A,B). The percentile bands around each condition mean show within-condition variability and inter - condition overlap .

**Figure 5.**
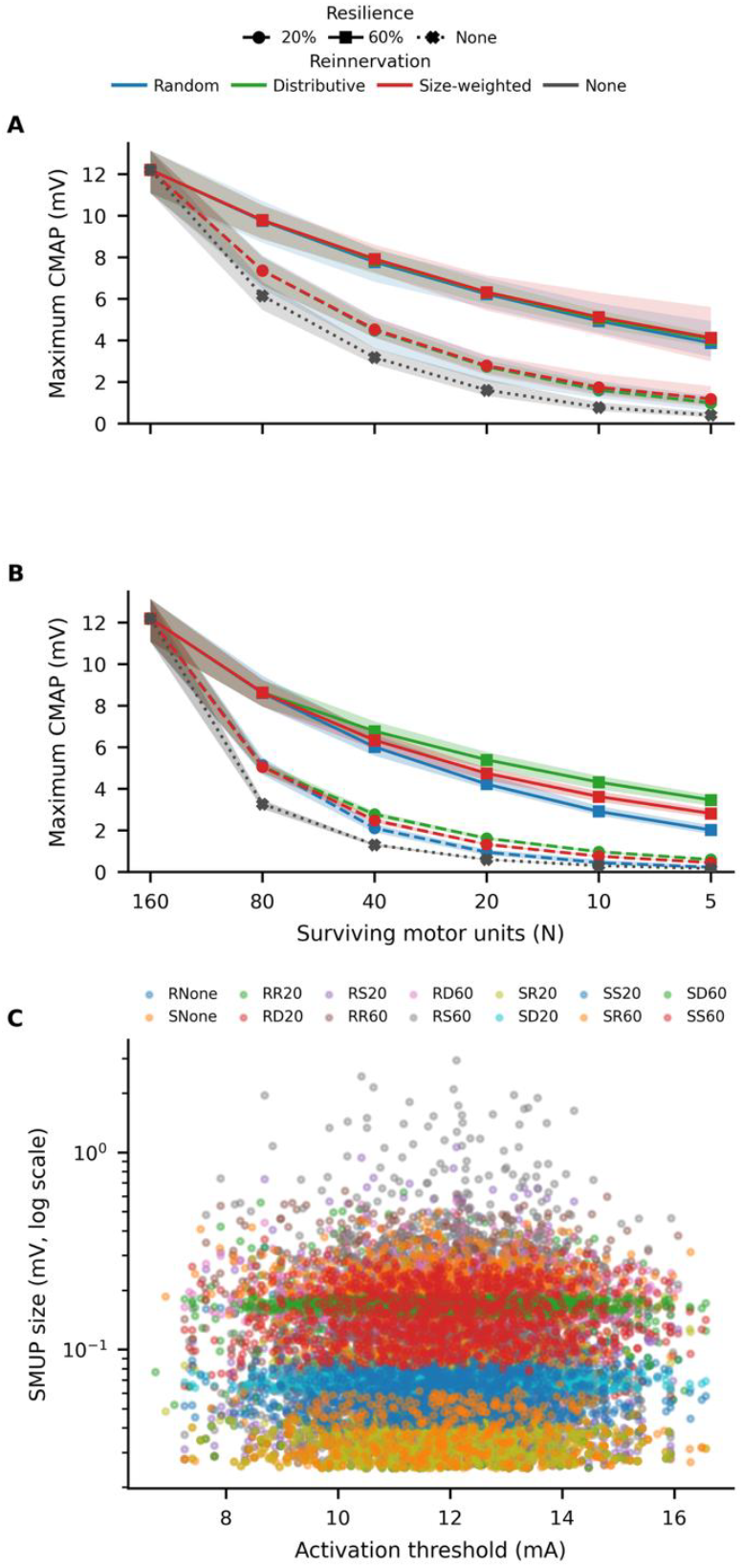
Simulated neuromuscular remodelling produced heterogeneous motor unit populations. (A,B) Maximum CMAP across progressive motor unit loss for conditions with random (A) or selective (B) denervation. Lines show condition means and shaded regions the 5th –95th percentile across 30 simulated trajectories. Colour identifies reinnervation mechanism, whereas line style and symbol identify resilience. (C) Distribution of motor unit activation thresholds and SMUP amplitudes at *N* = 40. Each point represents one motor unit from one of 30 trajectories for each of the 14 remodelling conditions; colours identify conditions. SMUP amplitude is shown on a logarithmic scale. The simulations produced within- and between-condition heterogeneity in both whole-muscle CMAP trajectories and the underlying motor unit populations.

To illustrate the underlying motor unit heterogeneity more directly, the distributions of activation thresholds and amplitudes at an intermediate stage of degeneration (*N* = 40) were examined (Figure 5C). Although activation thresholds remained broadly distributed over the initial starting range, SMUP amplitudes varied markedly across and within remodelling conditions, spanning more than two orders of magnitude. Thus, motor unit populations that could appear similar when summarized only by maximum CMAP were composed of substantially different combinations of unit size and threshold. Together these findings indicate that the simulations presented MScanFit with a diverse and physiologically heterogeneous set of motor unit populations.

### MOTOR UNIT NUMBER ESTIMATION ACCURACY

Across the 12 remodelling conditions that included collateral reinnervation (i.e., the 2 × 3 × 2 factorial design), MScanFit estimates followed the progressive reduction in true motor unit number across the 30 simulated trajectories within each condition (Figure 6). MScanFit underestimated the initial 160-unit pools to a similar extent across conditions, but mean estimates generally remained close to the true motor unit number as degeneration progressed. Differences between remodelling conditions became more apparent after motor unit loss and reinnervation began.

**Figure 6.**
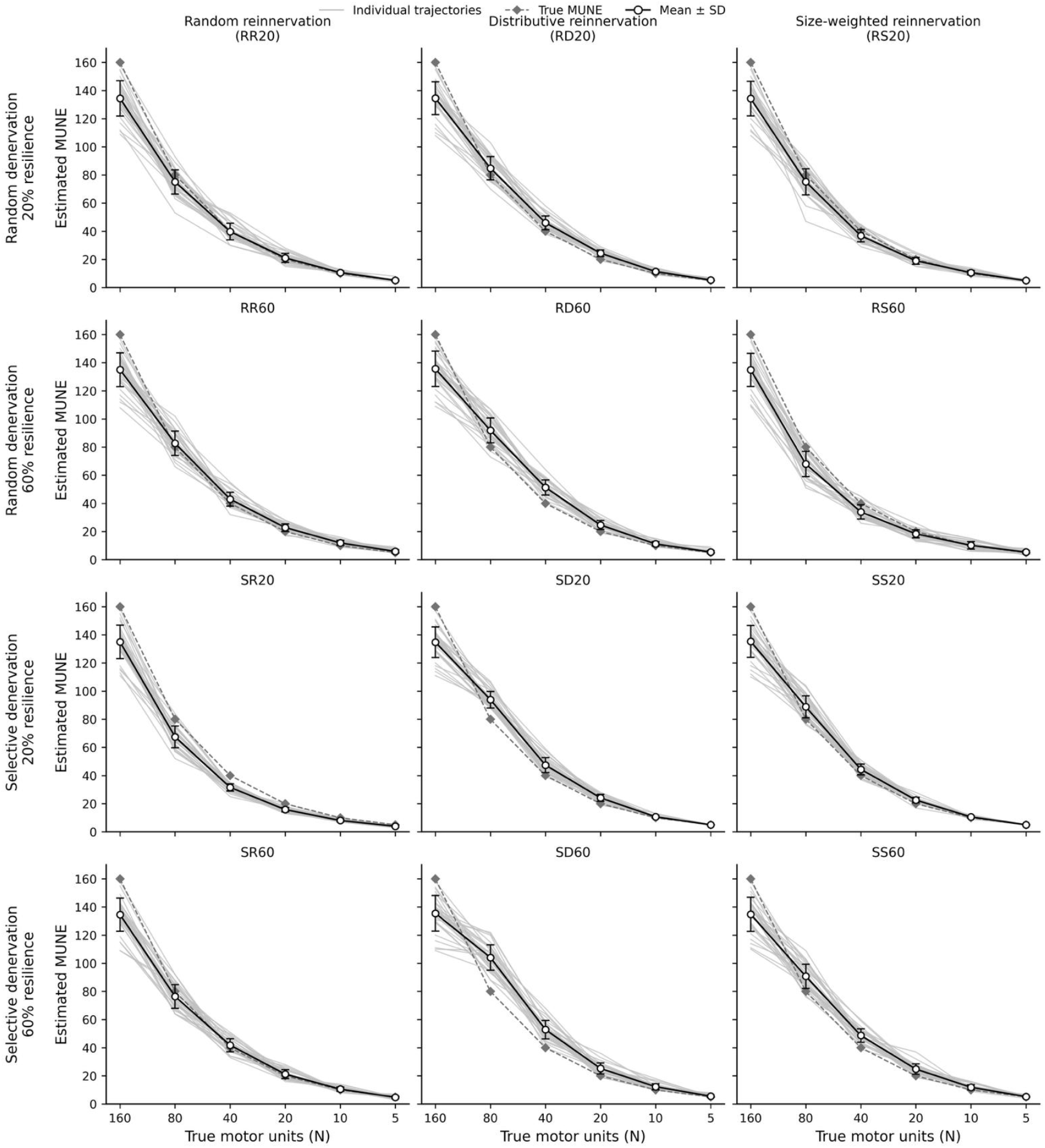
MScanFit MUNE estimates are broadly robust across simulated neuromuscular remodelling conditions. MScanFit continues to track motor unit loss as the underlying physiology changes over a disease-like trajectory. Each panel shows one condition from the primary 2 × 3 × 2 factorial design, arranged by denervation mechanism and resilience (rows) and reinnervation mechanism (columns). Light grey lines show the 30 simulated trajectories within each condition across true motor unit pool sizes of 160, 80, 40, 20, 10, and 5. Open circles wi th error bars indicate mean ± SD of the MUNE estimates at each stage. Dashed lines indicate the true number of motor units.

Signed MUNE error showed that the direction of estimation bias differed across remodelling conditions (Figure 7). Although MScanFit underestimated the 160-unit pools across all conditions, distinct error profiles emerged after remodelling began. For example, *RD60* showed predominantly positive errors, indicating overestimation of the true motor unit number, whereas *SR20* showed predominantly negative errors across the stages of motor unit loss. Thus, neuromuscular remodelling influenced not only the magnitude of MUNE error but also its direction.

**Figure 7.**
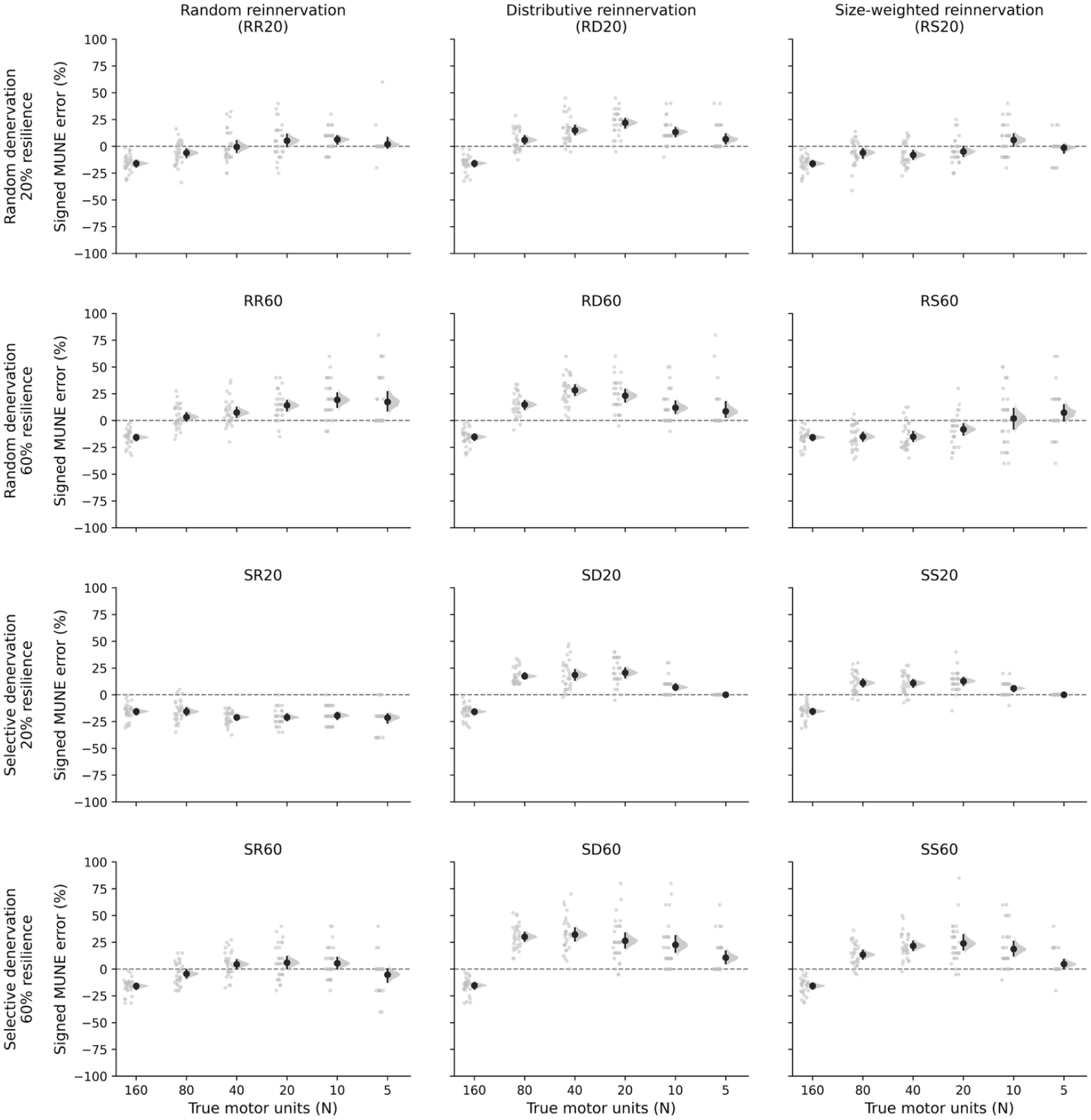
MScanFit MUNE is sensitive to the simulated neuromuscular remodelling conditions. Panels are arranged as in Figure 6. Instead of MUNE trajectories, each panel shows the signed MUNE percentage error at each stage of degeneration. Faint grey points represent the 30 simulated trajectories, black points indicate the mean signed error, vertical bars indicate 95% confidence intervals, and shaded half-violins show the bootstrap sampling distribution of themean error. Thehorizontal dashed line at 0% represents perfect agreement between the MScanFit estimate and the true motor unit number. Values below 0 indicate underestimation of the true motor unit number, whereas values above 0 indicate overestimation. The differing error profiles across panels indicate that MScanFit estimates are not invariant to the simulated remodelling physiology.

To determine whether the type of neuromuscular remodelling affected MScanFit accuracy, stage-specific marginal contrasts in MUNE absolute percentage error were estimated (Table 3). At 160 motor units, before remodelling, all contrasts were close to zero and their 95% confidence intervals included zero. Once remodelling began, selective denervation produced greater error than random denervation at 80, 40, and 20 motor units (mean differences of 5.0, 3.4, and 3.8 percentage points, respectively).

**Table 3.** Stage-specific effects of neuromuscular remodelling on MScanFit MUNE accuracy.

|  | Denervation | Reinnervation |  | Resilience |
|---|---|---|---|---|
| True MUs | Selective − Random | Distributive − Random | Size-weighted − Random | 60% − 20% |
| 160 | −0.1 [−1.6, 1.4] | −0.2 [−2.0, 1.6] | 0.0 [−1.9, 1.9] | −0.2 [−1.7, 1.3] |
| 80 | **5.0 [3.2, 6.8]** | **6.9 [4.9, 9.0]** | 1.6 [−0.5, 3.8] | **3.3 [1.5, 5.0]** |
| 40 | **3.4 [1.2, 5.5]** | **10.5 [7.7, 13.3]** | **2.6 [0.4, 4.8]** | **4.9 [2.7, 7.0]** |
| 20 | **3.8 [1.3, 6.3]** | **7.8 [4.7, 10.9]** | −0.3 [−2.8, 2.5] | **2.5 [0.1, 5.1]** |
| 10 | −1.0 [−3.7, 1.7] | −1.3 [−4.5, 2.1] | −0.3 [−3.3, 2.8] | **6.4 [3.8, 9.2]** |
| 5 | −1.3 [−4.2, 1.4] | **−6.7 [−10.3, −2.7]** | **−6.5 [−10.0, −3.0]** | **5.3 [2.6, 8.2]** |

*Values are marginal mean differences in MUNE absolute percentage error (percentage points) [95% CI]. Positive values indicate greater error for the first-named level. Bold values indicate 95% CIs that do not include zero. At each stage, n = 30 trajectories per condition (12 conditions; n = 360). The 160-MU stage precedes remodelling and serves as an internal control.*

Distributive reinnervation also produced greater error than random reinnervation at these stages, with the largest difference at 40 motor units (10.5 percentage points). Size - weighted reinnervation had smaller effects, with greater error than random reinne rvation at 40 motor units (2.6 percentage points), but lower error at 5 motor units (− 6.5 percentage points).

Resilience had the most consistent stage-specific effect. Resilience of 60% produced greater MUNE error at every stage after remodelling began, with mean differences ranging from 2.5 to 6.4 percentage points. Overall, the effects of different neuromuscular remodelling conditions were most apparent at intermediate stages of motor unit loss, whereas resilience affected MUNE accuracy across the full range of remodelled motor unit pools.

### MOTOR UNIT SIZE ESTIMATION ACCURACY

Across the 12 remodelling conditions, MScanFit estimates of mean single - unit amplitude generally followed the increase in target mean SMUP amplitude as motor unit number declined (Figure 8). Mean MSUE values remained close to their respective targets through much of the remodelling process, although variability among trajectories and differences between estimated and target amplitudes became more apparent as motor unit loss progressed, particularly in some of the 60% resilience conditions.

**Figure 8.**
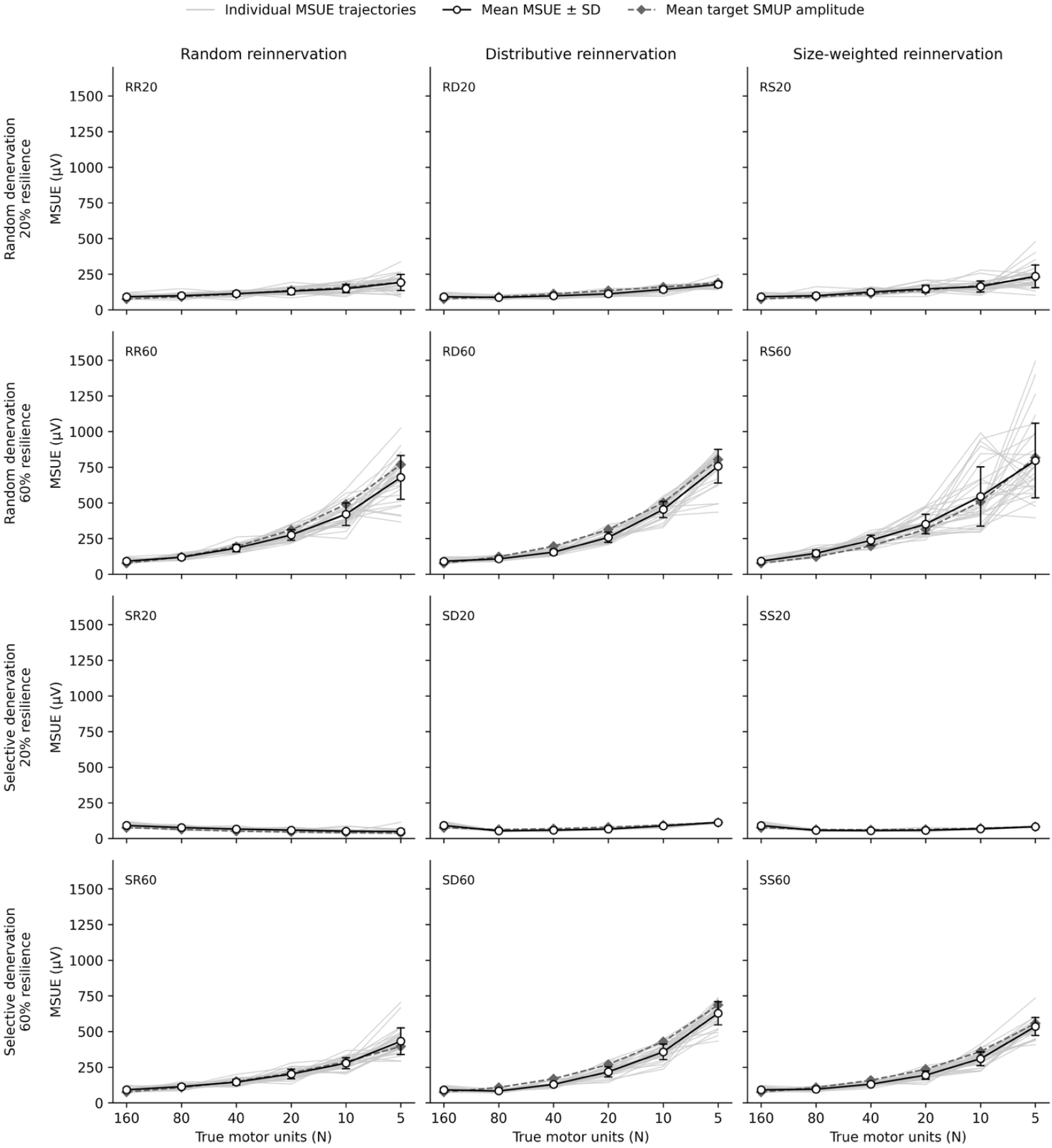
MScanFit MSUE across simulated neuromuscular remodelling conditions. Each panel shows one conditionfrom the primary 2 × 3 × 2 factorial design, arranged by denervation mechanism and resilience (rows) and reinnervation mechanism (columns). Light grey lines show the 30 simulated trajectories within each condition across true motor unit pool sizes of 160, 80, 40, 20, 10, and 5. Open circles wi th error bars indicate mean ± SD of the MSUE estimates at each stage. Dashed lines indicate the target mean SMUP amplitude of the corresponding simulated motor unit populations.

To determine whether the type of neuromuscular remodelling affected motor unit size estimation accuracy, stage-specific marginal contrasts in MSUE absolute error were estimated (Table 4). As with the MUNE analysis, prior to remodelling when all conditions started at 160 motor units, the contrasts were close to zero and the 95% confidence intervals included zero. After remodelling began, selective denervation generally produced lower error than random denervation, with differences emerging from 40 motor units and increasing at lower motor unit counts (− 5.7 *μ*V at 40 units to − 22.0 *μ*V at 5 units). Distributive reinnervation produced greater error than random reinnervation at 80, 40, and 20 motor units, with the largest difference at 20 units (12.9 *μ*V). Size-weighted reinnervation also produced greater error at several intermediate stages, including 40, 20, and 10 motor units. The largest and most consistent effect was associated with resilience: when compared with 20% resilience, 60% resilience increased MSUE absolute error at every stage after remodelling began, from 7.0 *μ*V at 80 motor units to 58.8 *μ*V at 5 motor units.

**Table 4.** Stage-specific effects of neuromuscular remodelling on MScanFit motor unit size estimation accuracy.

|  | Denervation | Reinnervation |  | Resilience |
|---|---|---|---|---|
| True MUs | Selective − Random | Distributive − Random | Size-weighted − Random | 60% − 20% |
| 160 | −0.2 [−2.1, 1.7] | −0.3 [−2.6, 2.0] | −0.1 [−2.4, 2.2] | −0.2 [−2.1, 1.7] |
| 80 | −0.5 [−2.5, 1.4] | **2.9 [0.9, 4.7]** | 2.3 [−0.3, 5.1] | **7.0 [4.9, 8.9]** |
| 40 | **−5.7 [−8.5, −3.1]** | **11.8 [8.8, 14.7]** | **8.8 [5.4, 12.1]** | **18.5 [15.8, 21.2]** |
| 20 | **−8.0 [−12.8, −3.5]** | **12.9 [8.0, 17.8]** | **5.4 [0.1, 11.5]** | **29.6 [25.2, 34.3]** |
| 10 | **−21.8 [−31.1, −13.4]** | 3.2 [−5.0, 12.2] | **16.0 [5.6, 27.9]** | **55.3 [46.9, 64.6]** |
| 5 | **−22.0 [−36.6, −8.9]** | −10.5 [−27.6, 6.0] | 1.0 [−17.0, 18.5] | **58.8 [45.8, 74.0]** |

*Values are marginal mean differences in MSUE absolute error (μV) [95% CI]. Positive values indicate greater error for the first-named level. Bold values indicate 95% CIs that do not include zero. At each stage, n = 30 trajectories per condition (12 conditions; n = 360). The 160-MU stage precedes remodelling and serves as an internal control.*

### ACCURACY IN THE ABSENCE OF COLLATERAL REINNERVATION

To isolate the impact of denervation alone, we included remodelling scenarios in the absence of collateral reinnervation. MScanFit accuracy depended strongly on the pattern of denervation (Figure 9). For MUNE, *RNone* and *SNone* showed similar absolute percentage error at 160 and 80 motor units but diverged markedly thereafter. Relative to *RNone, SNone* produced substantially greater MUNE error at 40, 20, 10, and 5 motor units, with mean differences growing from 16.5 to 30.7 percentage points. A similar pattern, with smaller magnitude, was observed for MSUE. Errors were comparable at 160 and 80 motor units, but *SNone* produced greater MSUE absolute error as more motor units were lost, with mean differences reaching 11.2 *μ*V at 5 motor units. Thus, when collateral sprouting was absent, selective denervation led to substantially poorer MScanFit accuracy than random denervation, particularly at advanced stages of motor unit loss.

**Figure 9.**
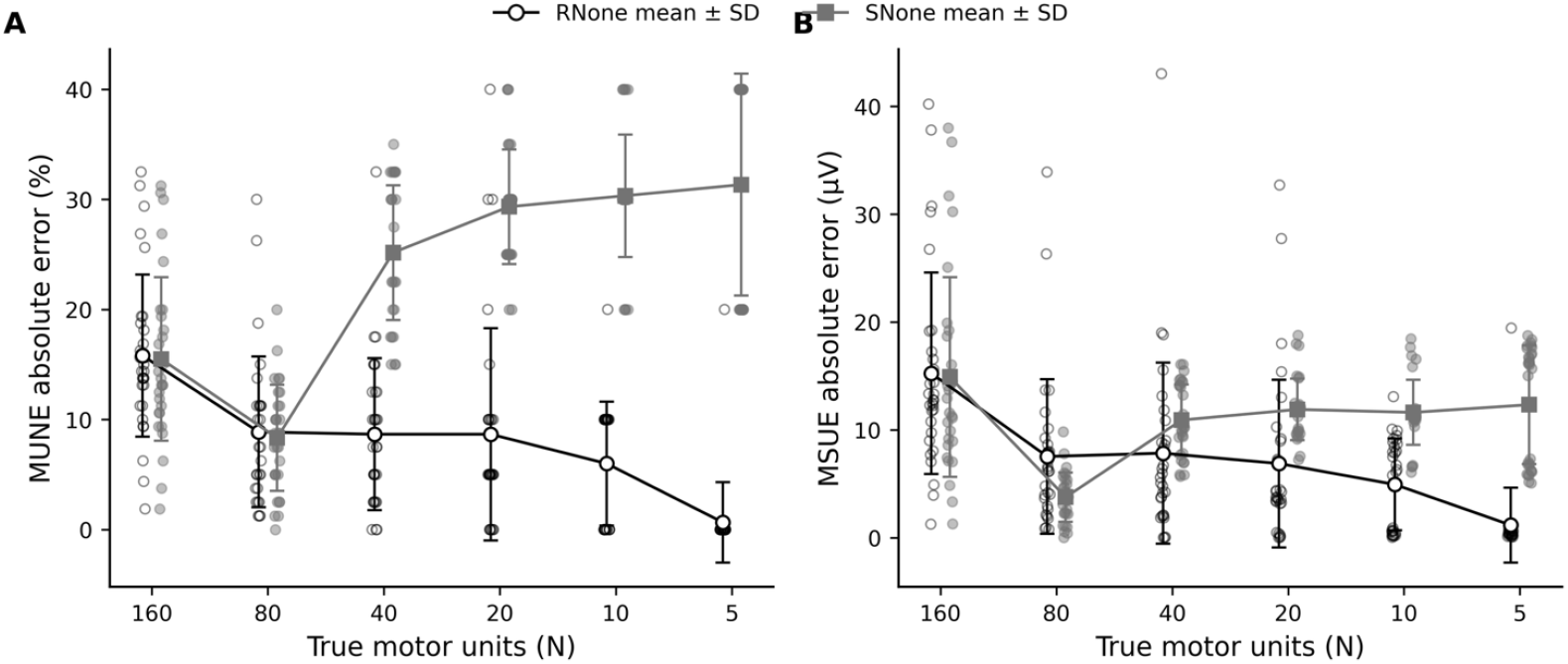
Effect of absent collateral reinnervation on MScanFit accuracy. Mean ± SD for (A) MUNE absolute percentage error and (B) MSUE absolute error across true motor unit pool sizes in the two no-reinnervation conditions, *RNone* and *SNone*. Individual simulated trajectories are shown as faint points. Points for the two conditions were offset slightly within each motor unit category to prevent overlap; the offset is for visual clarity only and has no analytical meaning.

## 4. Discussion

MScanFit estimates of motor unit number remained reasonably accurate across substantial motor unit loss and remodelling, supporting the conclusion that the algorithm is broadly robust to physiological heterogeneity (Figure 6). However, this robustness did not imply invariance: both the magnitude and direction of MUNE error changed systematically with the physiological rules used to generate the motor unit pool (Figure 7; Table 3). Selective denervation and distributive reinnervation produced stage-dependent changes in MUNE accuracy, whereas greater neuromuscular resilience produced the most consistent increase in error across remodelled motor unit pools (Table 3). Notably, remodelling conditions with similar maximum CMAP trajectories could nevertheless contain markedly different motor unit populations (Figure 5) and produce different patterns of MUNE bias (Figure 7). Motor unit size estimation was particularly sensitive to remodelling, with errors increasing with greater resilience and advanced motor unit loss (Figure 8; Table 4). Together, these findings indicate that MScanFit is comparatively robust for estimating motor unit number, while its errors remain dependent on the underlying neuromuscular remodelling process.

### BENCHMARKING MScanFit AGAINST THE ORIGINAL SIMULATION

Bostock introduced MScanFit as a practical method for estimating motor unit number and size from CMAP scans. 19 The concept of fitting the CMAP scan to infer the underlying motor unit population was not new; a Bayesian approach had previously been developed. 34,35 Bostock sought a more computationally tractable approach. As part of Bostock’s initial report, simulated CMAP scans with known motor unit populations were used to demonstrate the performance of the new fitting procedure across motor unit pools ranging from 160 to 5 units. The simulated degeneration model was deliberately simple: motor unit number was successively halved, with units removed randomly and some of the lost motor unit amplitude transferred to randomly selected surviving units to represent collateral reinnervation. Here, we reconstructed Bostock’s progressive motor unit loss and collateral reinnervation simulation as a benchmark for MScanFit (MSF1) and then systematically varied the physiological rules governing neuromuscular remodelling . Subsequent generative studies have used probabilistic CMAP models or alternative dynamic muscle models for algorithm development and biomarker evaluation, but addressed different questions. 23,26 We therefore retained Bostock’s six motor unit stages to provide a common benchmark before examining whether MScanFit performance remained robust when the assumptions governing neuromuscular remodelling were changed.

The *RR60* condition reproduced the principal behaviour reported by Bostock sufficiently well to establish this benchmark. In both studies, MSF1 estimates tracked the true decline in motor unit number across the full range from 160 to 5 units, although absolute percentage error was greater in *RR60*, particularly at the lower motor unit counts (Figure 4; Table 2). Overall MUNE absolute percentage error was 9.8 ± 10.3% in *RR60* compared with 6.9 ± 7.8% reported by Bostock, with the largest discrepancy at 5 –20 units. Exact numerical agreement was not expected because the two implementations differed in several respects (see Sections 2.4–2.6), but preservation of the principal relationship between true and estimated motor unit number provided a reference from which the effects of alternative remodelling physiology could be examined.

### MScanFit MUNE IS ROBUST, BUT NOT PHYSIOLOGICALLY INVARIANT

The primary objective of this study was to determine whether MScanFit remained accurate when motor unit loss occurred under different physiological rules for denervation and collateral reinnervation. The remodelling conditions produced different motor unit populations and whole-muscle CMAP trajectories (Figure 5), providing a substantive test of physiological robustness. Despite this heterogeneity, MScanFit continued to track the progressive reduction in true motor unit number across all 12 conditions with collateral reinnervation (Figure 6). Mean estimates generally remained close to the true motor unit number as the pools declined from 160 to 5 units. Thus, the principal finding is that MScanFit MUNE is broadly robust to substantial changes in the simulated physiology underlying motor unit loss.

This robustness did not, however, imply physiological invariance. The signed-error profiles in Figure 7 show that remodelling conditions developed distinct patterns of over - and underestimation once motor unit loss and reinnervation began. Particularly informative was the observation that conditions with similar maximum CMAP trajectories could nevertheless produce different MUNE error profiles. For example, the three 60% resilience conditions with random denervation showed considerable overlap in maximum CMAP across stages of motor unit loss (Figure 5A), yet their signed MUNE errors followed different trajectories (Figure 7). Similar preservation of the whole-muscle response therefore did not imply equivalent underlying motor unit populations (Figure 5C) or equivalent MScanFit bias. This distinction is physiologically relevant because collateral reinnervation can preserve CMAP amplitude despite substantial motor unit loss, as demonstrated in clinical observations of motor neuron disease and in dynamic muscle modelling.^36^ In the latter, an estimated 76 –91% of motor units could be lost before maximum CMAP fell below conventional lower limits.^26^ MScanFit uses information distributed across the complete CMAP scan, including its slope and variance, but its estimates necessarily depend on the motor unit sizes and activation thresholds that generate that scan. Bostock noted that unequal motor unit sizes alter CMAP variance and can bias the preliminary estimates used during MScanFit fitting.^19^ More recent simulation-based work has similarly shown that MUNE performance depends on properties of the generative CMAP model, including motor unit number and relative threshold spread, although that work was directed primarily towards development of an alternative estimator rather than the effects of progressive neuromuscular remodelling on MScanFit.^23^

The stage-specific contrasts quantify this lack of invariance (Table 3). Selective denervation and distributive reinnervation produced greater MUNE error predominantly at intermediate stages of motor unit loss, whereas size - weighted reinnervation produced smaller and less consistent effects. Greater neuromuscular resilience had the most consistent effect: 60% resilience produced greater absolute MUNE error than 20% resilience at every stage after remodelling began. Importantly, all contrasts were near zero at 160 motor units, before any remodelling had occurred, supporting the interpretation that these differences emerged from the imposed remodelling processes rather than differences in the initial motor unit populations. Even where differences were clear, however, the effect sizes were generally modest relative to the large changes in true motor unit number that MScanFit continued to track. The physiological factors examined here therefore modified MUNE error without negating the overall robustness of the estimate.

### COLLATERAL REINNERVATION AND MOTOR UNIT SIZE ESTIMATION

Motor unit size has long been recognized as a critical component of MUNE. Traditional MUNE methods estimate motor unit number from the ratio of the maximal muscle response to an estimate of mean surface motor unit potential size, making the result dependent on how representative the sampled motor units are of the underlying size distribution. Different approaches to calculating mean surface motor unit potential size can consequently produce substantially different MUNE values.^27^ Although MScanFit does not directly sample individual motor units in this way, it faces the related problem of inferring the number and size distribution of the motor units that generated the CMAP scan.

The full distribution of motor unit sizes may also be important. Using experimentally derived surface motor unit potential populations, van Dijk et al. showed that excluding small units altered MUNE substantially and could reduce sensitivity to changes associated with collateral reinnervation. 37 Thus, both the representation of mean motor unit size and the distribution from which that mean is derived have long been recognized as potential determinants of MUNE accuracy. The present simulations extend this issue to CMAP scan fitting by providing the true motor unit size distribution against which MScanFit estimates can be evaluated as that distribution is progressively altered by denervation and collateral reinnervation.

MScanFit generally followed the progressive increase in target mean SMUP amplitude as motor unit number declined, but the accuracy of mean unit-size estimates was strongly influenced by the remodelling conditions (Figure 8; Table 4). The clearest effect was associated with neuromuscular resilience. Compared with 20% resilience, 60% resilience increased MSUE absolute error at every stage after remodelling began, with the difference increasing from 7.0 *μ*V at 80 motor units to 58.8 *μ*V at 5 motor units. This effect follows directly from the model: greater resilience increased the extent of collateral reinnervation, and repeated remodelling progressively enlarged surviving units and broadened the distribution of SMUP amplitudes.

### DENERVATION WITHOUT REINNERVATION

The conditions without collateral reinnervation were included as boundary cases rather than as models of a typical course of motor neuron disease. Such boundary conditions are useful in generative models because they isolate the consequences of motor unit loss when compensatory remodelling is absent. This extreme is nevertheless biologically informative. In the SOD1G93A mouse, preferential loss of fast motor units can occur with limited enlargement of surviving fast motor units, whereas motor neurons in the same disease model can form substantially enlarged motor units following axonal regeneration.^8,11,38,39^ Thus, the capacity for compensatory reinnervation is not necessarily uniform across motor unit populations or physiological circumstances.

Removing collateral reinnervation in the present simulations exposed a marked dependence of MScanFit error on the denervation mechanism (Figure 9). Random and selective denervation produced similar errors early in degeneration, but diverged from 40 motor units onward. Selective denervation without collateral reinnervation produced progressively greater MUNE error, with the difference from random denervation increasing from 16.5 percentage points at 40 motor units to 30.7 percentage points at 5 motor units. The MSUE showed the same qualitative pattern, although the differences were smaller. Importantly, selective denervation did not have this effect in the factorial conditions where surviving units were allowed to remodel. The effect of denervation on MScanFit accuracy therefore depended on what happened to the surviving motor unit population after units were lost.

This result reinforces the broader conclusion that the physiological determinants of MScanFit error should not be interpreted independently. Denervation changes which motor units remain, whereas collateral reinnervation subsequently changes their sizes and contribution to the CMAP scan. The motor unit population presented to MScanFit is therefore the product of both processes, and different combinations can generate emergent patterns of estimation error.

### LIMITATIONS

#### Biological simplification

Our generative model operates at the level of SMUP amplitude and activation threshold rather than individual muscle fibres and complete motor unit potential waveforms. It therefore does not explicitly represent fibre spatial distribution, waveform morphology, temporal dispersion, phase cancellation, conduction velocity heterogeneity, or disease-related changes in single-fibre action potentials, all of which can influence surface EMG and CMAP morphology.^40–42^ Bostock’s original MScanFit simulation similarly assumed that each motor unit contributed a constant increment to peak CMAP amplitude and identified this as an important simplification. 19 Selective vulnerability was also deliberately simplified: selective denervation was implemented by preferential removal of motor units with the largest current SMUP amplitudes. This captures one plausible feature of motor neuron disease but does not explic itly model slow, fast fatigue-resistant, and fast fatigable motor unit subtypes or the mechanisms underlying their differential vulnerability.^2,7,9–11,38,39^ Similarly, the random, distributive, and size-weighted reinnervation rules were designed to span a physiologically plausible remodelling space rather than to represent mutually exclusive mechanisms of reinnervation in vivo.

#### Model and experimental simplification

Neuromuscular resilience was defined operationally as a model parameter controlling the extent of collateral reinnervation and was examined at 20% and 60%. These values represent controlled differences in modelled collateral reinnervation efficacy and should not be interpreted as established biological categories or directly measurable patient-level resilience metrics, although experimental and modelling studies support substantial variation in reinnervation capacity in motor neuron disease.^11,43^ In experimental models of extensive partial denervation (up to 85–90% motor unit loss), surviving motor units expand 2 - to 3-fold under stable conditions up to an upper physiological ceiling of 4 - to 6-fold.^44^ In our sequential halving framework, resilience levels of 20% and 60% yield cumulative mean unit enlargements of approximately 1.7-fold and 4.1-fold, respectively, at 87.5% motor unit loss (N = 20). The degeneration stages (160→80→40→ 20→ 10→5 units) were retained from Bostock to facilitate benchmarking and represent increasing severity of motor unit loss rather than equal intervals of disease progression or a proposed temporal trajectory of ALS.

Activation physiology was also simplified: SMUP amplitude and activation threshold were sampled independently, threshold variability was constrained to a fixed relative spread, and disease-related changes in axonal excitability or threshold heterogeneity were not modelled .

Several experimental sources of variability were intentionally absent. Electrode placement, stimulation geometry, recording-system characteristics, skin–electrode impedance, participant positioning, movement artefacts, and biological variation between visits were not represented, although multicentre MUNE and MUNIX studies have shown that recording conditions and operator-dependent variation in CMAP amplitude can affect measurement stability.^22,45–47^ The present study therefore evaluates MScanFit accuracy under controlled simulated physiology, not clinical reliability or test –retest repeatability. Implementation choices may also influence the results. We used the default MScanFit fitting procedure (MSF2), a minimum motor unit amplitude of 25 *μ*V, and prescribed pre- and post-scan regions; the simulated SMUP distribution was bounded at the same 25-*μ*V minimum, so smaller units were absent by design. Bostock varied the minimum unit - size constraint according to estimated unit size, and subsequent multicentre work highlighted sensitivity of MScanFit to CMAP amplitude and analysis settings, contributing to later modifications of the fitting procedure.^47^ Version-specific settings and recording conditions may therefore contribute to differences between the present simulations and experimental MScanFit studies.

#### Generalizability

Model calibration was based on CMAP scans from the abductor pollicis brevis (APB), and the initial motor unit population and SMUP amplitude distribution were chosen to reproduce characteristics of experimental APB scans.^22^ The results therefore apply most directly to APB-like recordings and should not be assumed to generalize to muscles with substantially different architecture, innervation, or CMAP scan characteristics. More broadly, surface-recorded MUNE and MUNIX sample only the portion of the motor unit population represented by the recording configuration, and estimates are known to be both method- and muscle-dependent.^41,48–50^ The present APB model therefore provides a controlled framework for isolating the effects of neuromuscular remodelling on MScanFit, but further work is required to determine whether the same dependencies occur in other muscles, recording configurations, and CMAP-scan-based methods such as STEPIX, CDIX, and StairFit.^15–17^

## 5. Conclusion

MScanFit estimates of motor unit number remained reasonably accurate across substantial motor unit loss and heterogeneous neuromuscular remodelling, demonstrating broad physiological robustness. This robustness did not imply invariance, as both the magnitude and direction of MUNE error depended systematically on the processes governing denervation and collateral reinnervation. Motor unit size estimat es were also strongly influenced by remodelling, particularly as collateral reinnervation increased and motor unit pools became small. These findings support MScanFit as a robust approach for estimating motor unit loss across heterogeneous neuromuscular phenotypes and indicate that some variability in its estimates reflects underlying physiological differences in the motor unit population.

## Data availability

The simulated CMAP scans, corresponding MScanFit outputs, and analysis-ready dataset are available on Figshare at https://doi.org/10.6084/m9.figshare.33190827

## Author contributions

DA: methodology; software; formal analysis; data curation; visualization; writing (original draft)

MA: conceptualization; methodology; software; writing (review & editing)

KEJ: conceptualization; methodology; supervision; formal analysis; validation; visualization; writing (review & editing)

All authors approved the final version of the manuscript and agree to be accountable for all aspects of the work.

## Declaration of generative AI and AI-assisted technologies in manuscript preparation

During preparation of this manuscript, the authors used ChatGPT (OpenAI) and Gemini (Google) to assist with organization, language refinement, and critical review of manuscript text. All scientific content, analyses, interpretations, citations, and revisions were reviewed and verified by the authors, who take full responsibility for the final manuscript.

